# Allometric scaling of RNA abundance from genes to communities

**DOI:** 10.1101/2021.10.03.462954

**Authors:** Mark Louie D. Lopez, Ya-ying Lin, Stephan Q. Schneider, Ryuji J. Machida

## Abstract

The metabolic theory of ecology (MTE) and growth rate hypothesis (GRH) help explain the mechanistic basis of size (allometry) and temperature dependence on growth rate and whole-body-RNA content in organisms. However, testing RNA allometric scaling with next-generation sequencing is yet to be done. Here, we validated the assumptions of GRH and MTE on messenger RNA and ribosome abundance using mock community metatranscriptome analysis. Our findings highlight that fast-growing smaller species harbor greater RNA abundance per mass of tissue compared with species having larger body sizes and slower growth rates, where allometric slopes for genomic and gene-level RNA abundance range from –⅓ to −1. We found that genome size and body size impose significant constraints in interspecific RNA abundance scaling, while the assumed temperature dependence appeared to be weak. Lastly, allometric scaling integration in community-level models may extend the use of metatranscriptomics as a reliable tool for estimating ecosystem processes.

## Introduction

Understanding the mechanistic basis of individual metabolism and growth has been one of the core questions in ecology. These affect energy flux, nutrient storage, and the overall turnover of materials in the ecosystems. The metabolic theory of ecology (MTE) and the growth rate hypothesis (GRH) have significantly advanced our understanding of the allometric scaling of these processes (1, 2). Allometry describes the changes in physiological rates in organisms due to proportional changes in body size (3). While the MTE explains the importance of energy as the primary driving factor for metabolism (4), GRH deals with the dynamic patterns of energy and primary elements in living systems (5).

Specifically, MTE suggests that the metabolic rate of organisms changes proportionately with body size and temperature. With the fluid dynamics principle, MTE proposed a universal ¾ power scaling for metabolic rate (–¼ for mass-specific metabolic rate) that describes the optimal supply rate towards different cells through the transport networks in organisms. Moreover, it assumes that metabolic rate tends to be higher in individuals at a warmer temperature than those of the same size living in a colder environment (Fig. 1A). On the other hand, GRH expects smaller and fast-growing species to require higher phosphorus (P) allocation and increased demand for P-rich ribosomal RNA needed to sustain higher growth rates (6). The GRH also offers a genetic approach to explaining natural selection on genome size and growth rate (Fig. 1B), where studies have demonstrated that increased growth rate and transcriptional capacity of ribosomal RNA production are positively associated with gene length and genome size due to the presence of multiple copies of intergenic spacers (5, 7).

**Fig. 1.**
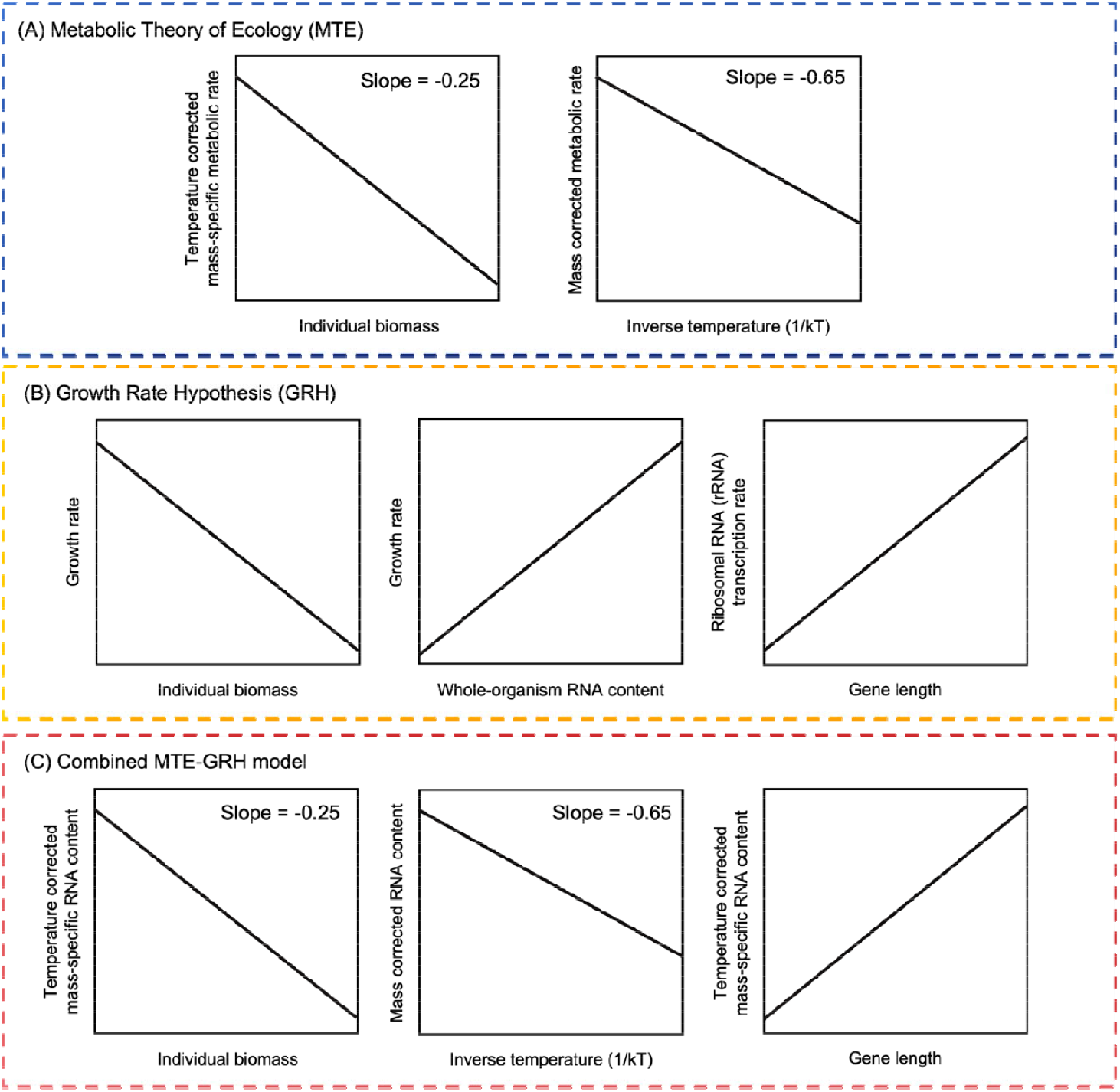
Assumptions of the validated principles in the study: (A) metabolic theory of ecology (MTE), (B) growth rate hypothesis (GRH), and the extended MTE-GRH model in demonstrating how body size, gene length, and temperature constrain growth rate and RNA content from genes to the ecosystem level.

In relating individual metabolic rate with growth, MTE assumptions were extended to link whole-body P and RNA content with the energy fluxes in organisms (MTE-GRH: Fig. 1C). The MTE-GRH predicts that the whole-body P and RNA content are tightly linked to energy flux in organisms based on mass- and temperature-dependence of ATP production in mitochondria that should influence growth rate (8). Combining these ideas, both MTE and GRH demonstrate how body size and temperature constrain growth rate, RNA content (e.g., ribosomal concentration), and elemental composition from genes to the ecosystem level (9). The correlation between two biological currencies: energy through ATP and materials in the form of phosphorus-rich RNA are relevant in understanding the ecosystem functions. Although both assumptions of MTE and GRH have been suggested as plausible mechanisms for regulating the RNA metabolic process (8), the generality of these arguments awaits genomic and gene-level empirical evidence using next-generation sequencing (NGS) technologies.

Here, we tried to validate the assumptions of GRH and MTE on allometric scaling of messenger RNA (mRNA) transcript and ribosome, with ribosomal RNA (rRNA) as a proxy, abundance using metatranscriptome analysis of model organism mock communities. Metatranscriptomics is a sequencing tool that uses NGS to detect and quantify RNA in a community sample at a given time point (10). Based on the MTE-GRH model (8), transcription rate (TR), which refers to mRNA and rRNA production per unit of time, is expected to vary with body size and temperature:

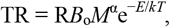

where R is the transcript read number expended per unit mass of tissue (read numbers/g), *B_o_* is the normalization coefficient, *M* is the individual body mass (in g), α is the allometric coefficient, and e^−*E*/*kT*^ is the Arrhenius factor reflecting the temperature dependence of metabolic rate, where *E* is the average metabolic activation energy (0.65 eV), *k* is Boltzmann’s constant (8.62 × 10^−5^ eV K^−1^), and *T* is the absolute temperature in kelvins (11). Rearranging these terms from the equation and taking the natural logarithms (ln) yields: ln(TRe^*E/kT*^) = αln*M* + *C*, where *C* = ln(R*B*_o_). This model predicts that plotting the ln of temperature-corrected transcript abundance (ln[Re^*E*/*kT*^] (read number/g)) should give a linear function of ln(*M*) with a slope or allometric coefficient (α) of approximately −¼ based on its origins in the fractal-like geometry of biological exchange surfaces and distribution networks (4). At the same time, ln of mass-corrected transcript abundance ([ln(R*M*^α^] (read numbers/g)) should be a linear function of the inverse of temperature (1/*kT*) with a slope of negative activation energy value (−*E or* −0.65 eV), reflecting the kinetics of aerobic metabolic processes in organisms (4, 8, 11). Lastly, the RNA abundance per mass of tissue should be positively associated with gene length and genome size due to the presence of multiple copies of intergenic spacers (5, 7).

To validate the model, we constructed mock communities consisting of five species spanning wide orders of magnitude in body size incubated at three temperature conditions. Using the mock communities, thorough genome-wide analysis on the power-scaling of interspecific RNA abundance is possible using selected model species with the available reference genome sequence. At the genomic level, overall nuclear and mitochondrial-encoded mRNA transcripts and ribosome abundance were measured. Moreover, power-scaling for selected mitochondrial and nuclear-encoded mRNA and rRNA genes was analyzed. The results from this study could provide a unifying explanation of varying RNA transcription patterns among different species and expand the use of metatranscriptomics in estimating active biological processes (e.g., growth rate and biomass) for community ecology studies.

## Results

### Mock community analysis of RNA abundance

Using prepared RNA libraries from pooled homogenized individuals of incubated model organisms (Fig. S1), we successfully accounted for the proportional difference in RNA abundance among species with a wide range of body and genome sizes. The significant differences in the body (1 to 40,000 μg) and genome (C-values, 0.11 to 1.75 pg) masses and the availability of high-quality reference genome sequences for each species provided a perfect premise to see the effect of these variables on the interspecific allometric scaling of RNA abundance in community samples.

### Genomic-level empirical model validation

In support of the assumption of MTE, the logarithmic relationships of temperature-corrected transcript abundance (Re^*E*/*kT*^) and individual body mass are in negative linear correlation for all transcripts (Fig. 2A). This shows that faster-growing smaller species harbor more mRNA transcripts and ribosomes (deduced from rRNA transcript reads) per tissue mass than species with larger body sizes and slower growth rates. This observation supports the predictions of the MTE-GRH model concerning the relationship between RNA abundance, growth rate, and body size of organisms. In terms of the slope, only the mitochondrial rRNA among the four transcripts has 95% confidence intervals that include the value predicted by MTE of −0.25 (Table 1). The three other cellular components: mitochondrial mRNA, nuclear mRNA, and nuclear-encoded ribosomal RNA, exhibited a steeper slope closer to −0.33. It can be noted that the nuclear-encoded mRNA and ribosome have steeper slopes (−0.40 and −0.33) than the mitochondrial-encoded counterparts (−0.30 and −0.28). All four fitted lines in Fig. 2A explain 51–86% (adjusted *R*^2^, Table 1) of the variation in ln-transformed transcript read abundance among species. However, mRNA transcripts and ribosome abundance are observed to be less sensitive to the changes in temperature (Fig. 2B), where the fitted slopes range from −0.02 to −0.13, which significantly differ from the slope predicted by MTE (−0.65, Table 1). The mass-specific transcript abundance for the species peaks at an optimal temperature for growth of 19 °C, dropping as the temperature increases to 28 °C (Fig. S2).

**Fig. 2.**
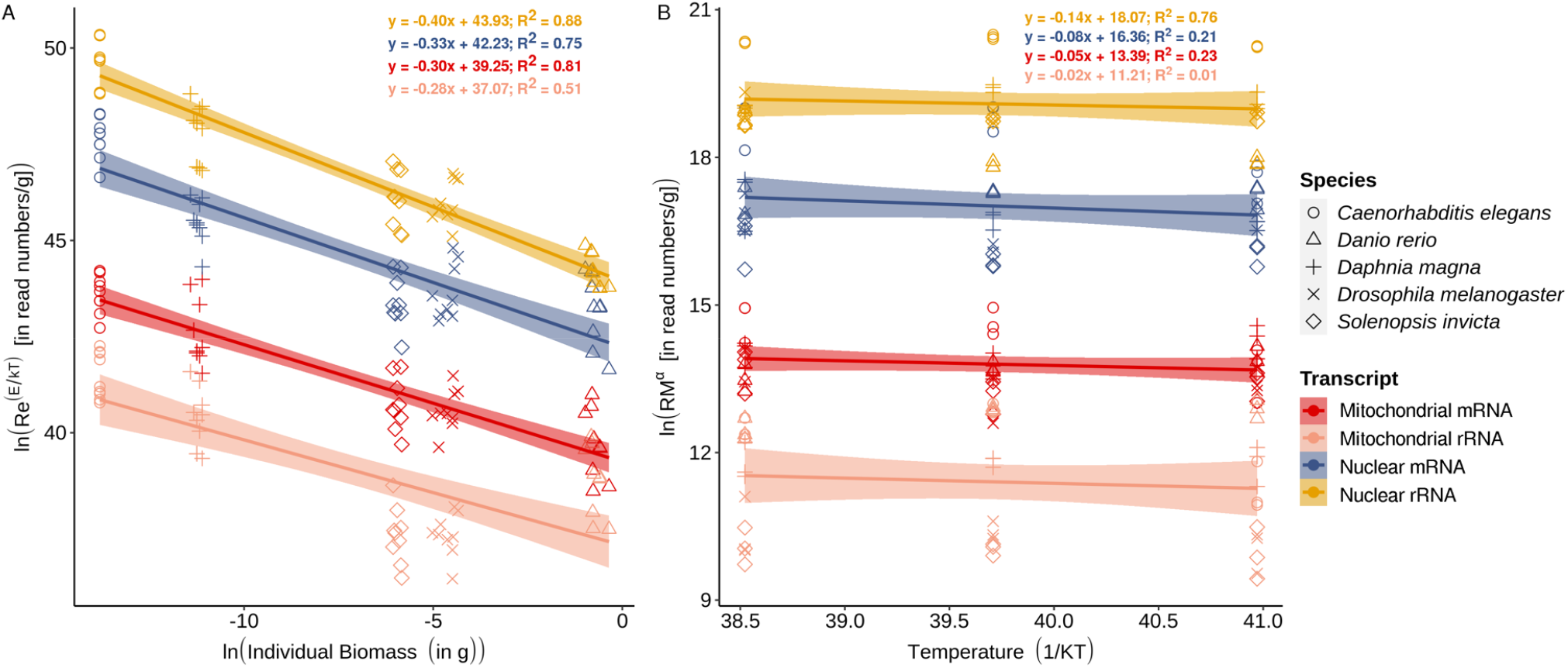
Effect of (A) individual body mass (in g) on temperature-corrected transcript abundance ([Re^*E*/*kT*^] (in read numbers/g)) and (B) the effect of temperature (1/*kT*) on mass-corrected transcript abundance (*RM*^α^ [in read numbers/g]) of species from the constructed mock communities. For the variable: R refers to transcript read count (read numbers/g); e^*E*/*kT*^ is the Arrhenius factor, where *E* is the average activation energy for metabolic processes (0.65 eV), *k* is Boltzmann’s constant (8.62 × 10^−5^ eV K^−1^), and *T* is the absolute temperature in kelvins; *M*^α^ is the body size term, *M*^1/4^ is based on the fractal-like geometry of biological exchange surfaces according to the assumptions of MTE.

**Table 1.** Parameter estimates for the regressions depicted in Fig. 2A [ln(Re^*E*/*kT*^) vs ln(*M*)] and Fig. 2B [ln(R*M*^1/4^) vs 1/*kT*]. Bold texts reflect significant correlation between variables.

| Transcript | $\ln(\text{Re}^{E/kT})$ vs $\ln(M)$ | | | | | $\ln(RM^{1/4})$ vs $1/kT$ | | | | |
| --- | --- | --- | --- | --- | --- | --- | --- | --- | --- | --- |
| | Fitted slope ± | Predicted | Intercept | Adjusted | *P*- | Fitted slope ± | Predicted | Intercept | Adjusted | *P*- |
| | 95% CI | slope | | $R^2$ | value | 95% CI | slope | | $R^2$ | value |
| Mitochondrial mRNA | $-0.30 \pm 0.02$ | -0.25 | 39.25 | 0.80 | **<0.01** | $-0.05 \pm 0.01$ | -0.65 | 13.38 | 0.23 | **<0.01** |
| Mitochondrial rRNA | $-0.28 \pm 0.04$ | -0.25 | 37.06 | 0.51 | **<0.01** | $-0.02 \pm 0.03$ | -0.65 | 11.21 | 0.01 | **<0.01** |
| Nuclear mRNA | $-0.33 \pm 0.03$ | -0.25 | 42.22 | 0.75 | **<0.01** | $-0.08 \pm 0.02$ | -0.65 | 16.36 | 0.20 | >0.01 |
| Nuclear rRNA | $-0.40 \pm 0.02$ | -0.25 | 43.93 | 0.88 | **<0.01** | $-0.14 \pm 0.01$ | -0.65 | 18.07 | 0.76 | **<0.01** |

### Gene-level empirical model validation

At the gene level, the linear relationships between ln-transformed temperature-corrected transcript abundance (Re^*E*/*kT*^) and individual body mass for 10 mitochondrial protein-coding, two mitochondrial rRNA (small [*12S*] and large [*16S* ribosomal RNA), and two nuclear rRNA (small [*18S*] and large [*26S* or *28S*] ribosomal RNA) genes (Fig. 3A) are all negatively correlated, which shows a similar pattern to the genomic transcripts. Again, this provides consistent evidence for the assumptions of the MTE-GRH model that gene-level mRNA transcript abundance per gram of tissue must be higher in smaller fast-growing species than larger ones. However, the fitted slope for most genes is steeper than the slope predicted by MTE (−0.25). The average fitted slope for the selected mitochondrial genes and nuclear-encoded rRNA are −0.30 and −0.47, respectively. Like genomic-level observation, nuclear-encoded genes tend to have steeper slopes than mitochondrial genes. The fitted lines for the selected mitochondrial and nuclear-encoded genes explain 43–86% of the variation in gene-level transcript abundance (adjusted *R*^2^ Table S3). In assessing whether metabolic scaling can improve the predicted model between transcript read abundance and species biomass, temperature and body size scaling were compared in this study. Table 2 shows that accounting for the effect of body size using the allometric coefficient for the selected gene markers (ln[R] vs ln[*M*^α^]) can improve the *R*^2^ value from around 0.01 to 0.06 compared with uncorrected values that exclude temperature and body size effects. Meanwhile, temperature-corrected scaling provided the lowest *R*^2^ values among the predicted models.

**Fig. 3.**
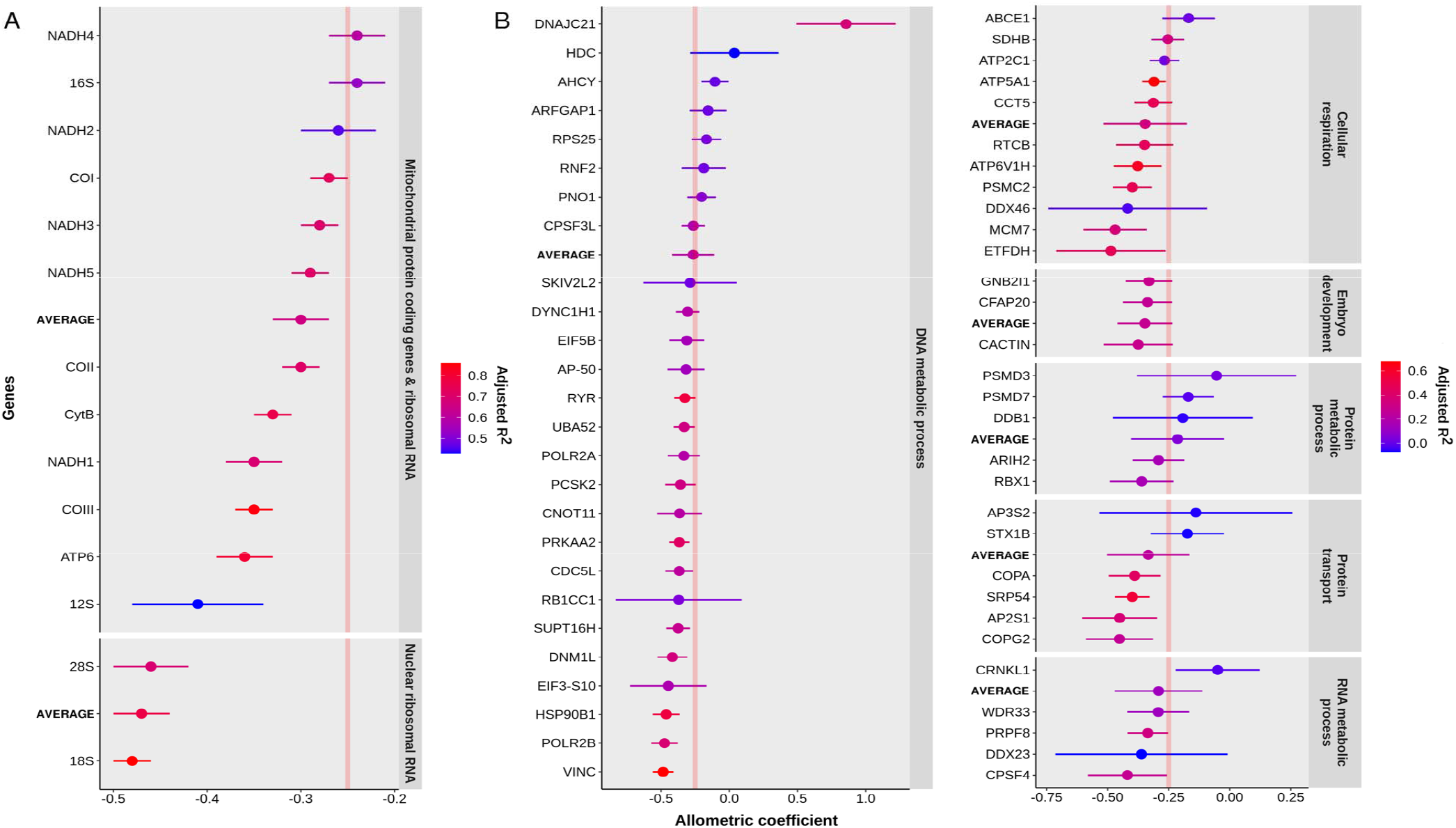
Allometric coefficients (±95% confidence interval) for (A) selected nuclear and mitochondrial genes and (B) annotated nuclear-encoded orthologous genes based on the natural log relationship of temperature-corrected transcript abundance ([Re^*E*/*kT*^] (in TPM/g)) and individual body mass (in g) linear regression. The red line indicates the predicted allometric coefficient according to MTE (−0.25).

**Table 2.** Comparison of the adjusted *R* (*R*^2^) for the correlation between: uncorrected masss-pecific transcript abundance (R [reads/g]) and individual body mass (g), temperature-corrected transcript abundance ([Re^*E*/*kT*^] (in read numbers/g)) and individual body mass, and mass-specific transcript abundance (R [reads/g]) and allometrically scaled biomass (*M*^α^ [in g]).

| Source | Genes | $\ln(R)$ vs $\ln(M)$ | $\ln(Re^{E/kT})$ vs $\ln(M)$ | $\ln(R)$ vs $\ln(M^a)$ |
| --- | --- | --- | --- | --- |
| Nuclear | *18S* | 0.92 | 0.86 | 0.96 |
|  | *28S* | 0.73 | 0.69 | 0.79 |
| Mitochondrial | *12S* | 0.44 | 0.43 | 0.45 |
|  | *16S* | 0.59 | 0.53 | 0.62 |
|  | COI | 0.82 | 0.73 | 0.87 |
|  | CytB | 0.84 | 0.76 | 0.90 |

Moreover, 62 nuclear-encoded orthologous genes with enough transcript reads for each species in the mock community were classified into six Gene Ontology (GO) terms (Fig. 3B). The ln-relationships of the temperature-corrected transcript abundance ([Re^*E*/*kT*^ (in transcripts per million/g)) and individual body mass was predominantly negatively correlated [except *DNAJ homolog*, *subfamily C*, *member 21* (DNAJ21), and *Histidine decarboxylase* (HDC) genes] among the examined genes. The fitted slopes for the genes range from −0.48 to 0.85, where some genes have an allometric coefficient close to the −0.25 assumption of MTE (red line, Fig. 3B). Moreover, the average slope for the six identified GO terms reflected comparable values to the MTE-predicted slope (from −0.34 to −0.21). Lastly, the fitted lines for the nuclear-encoded orthologous genes only explain 1–54% of the variation in gene-level transcript abundance (adjusted *R*^2^, Table S5), which shows that the body size has a weak effect on the transcription process of individual genes.

### Genome size and RNA scaling

Despite the observed mass dependence of mRNA transcript and ribosome abundance, considerable residual variations probably indicate the importance of other factors aside from body size. We validated the genetic basis of GRH that correlates RNA transcription and genome size by examining the ln-relationship of temperature-corrected transcript abundance (Re^*E*/*kT*^) and genome size (Fig. 4). A negative linear correlation was observed between genome size and transcript read abundance. This indicates that species with smaller genome sizes exhibit a more significant number of transcriptomic reads per gram of tissue than species with more DNA content in the nucleus. The fitted lines explain 16–67% of the variation in mRNA transcript and ribosome abundance. Overall, individual body size, the genome size (DNA content), and their interaction explained a substantial percentage (70%) of the variance in transcript read abundance among species (Table 3). This implies that aside from body size, the genome size of the species also imposes constraints on transcriptomic metabolism.

**Fig. 4.**
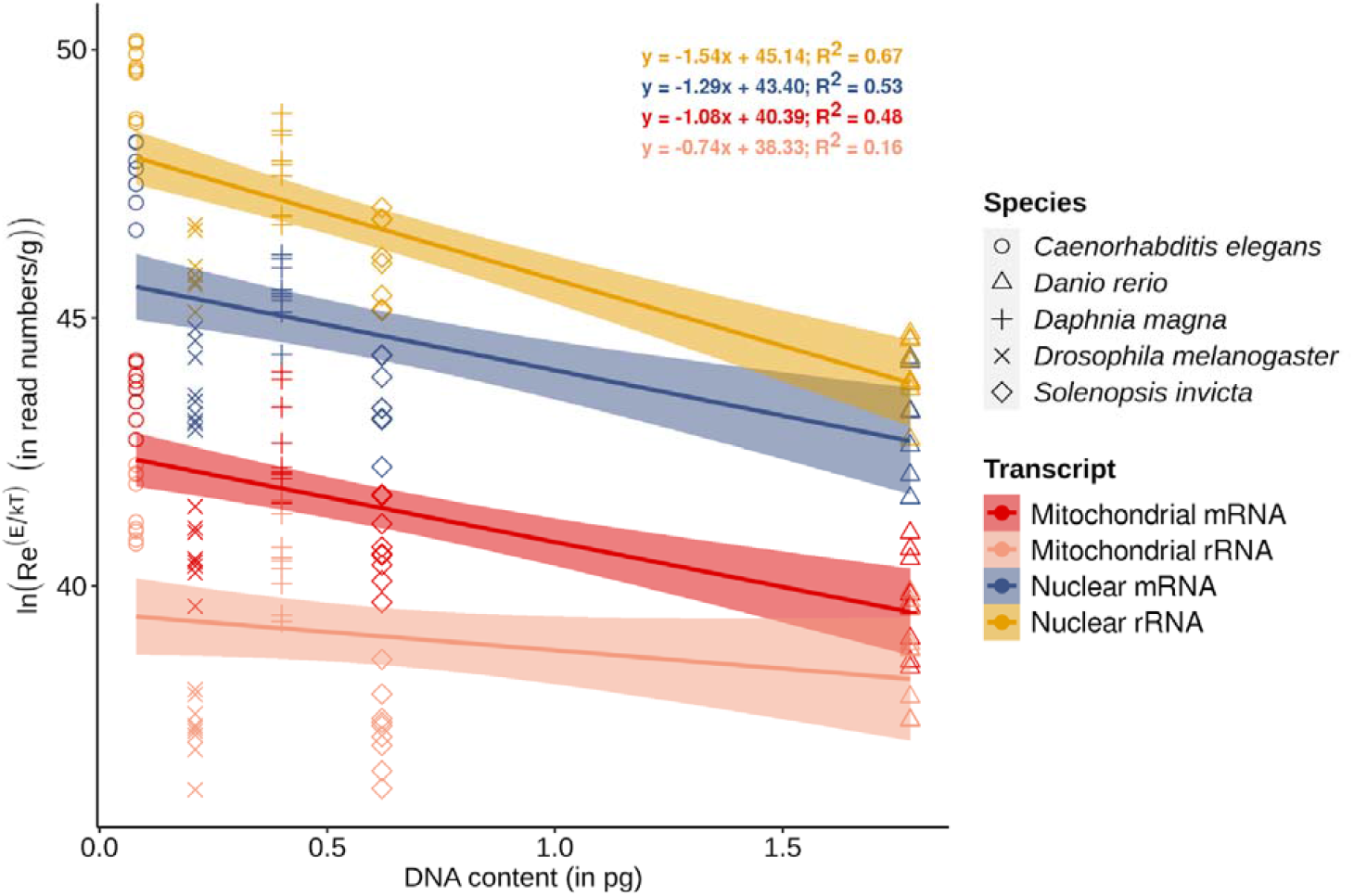
Effect of genome size [DNA content (in pg)] on temperature-corrected transcript abundance ([Re^*E*/*kT*^] (in read numbers/g)) of species from the constructed mock communities. For the variables: R refers to mass-specific transcript read count (read number/g), and e^*E*/*kT*^ is the Arrhenius factor, where *E* is the average activation energy for metabolic processes (0.65 eV) and *k* is Boltzmann’s constant (8.62 × 10^−5^ eV K^−1^).

**Table 3.** Results of ANCOVA to analyze individual body mass, DNA content, and temperature, and their interactive effects as covariates in temperature-corrected transcript abundance. Bold texts reflect the significant contribution of the covariant.

| Source of variation | df | *F* | *P*-value | Effect size |
| --- | --- | --- | --- | --- |
| Individual body mass | 1 | 50.94 | **<0.01** | 0.23 |
| DNA content | 1 | 161.45 | **<0.01** | 0.48 |
| Temperature | 1 | 0.16 | 0.682 | 9.99 ×10<sup>-4</sup> |
| Individual body mass: Temperature | 1 | 4.40 | **<0.01** | 0.02 |
| DNA content: Temperature | 1 | 0.31 | 0.575 | 2.00 × 10<sup>-3</sup> |
| Individual body mass: DNA content | 1 | 400.52 | **<0.01** | 0.70 |
| Individual body mass: DNA content:<br>Temperature | 1 | 5.30 | **<0.01** | 0.03 |

Interestingly, we found a negative correlation between gene length and the allometric coefficient of the annotated genes (Fig. 5). This means that transcript read abundance of longer genes is more sensitive to changes in body size. Although the coefficients of determination are low, the observation of negative slope was consistent throughout all GO terms. Overall, these observations provide supporting evidence on the genetic basis for the relationship between RNA metabolism, growth, and body size according to the assumptions of the MTE and GRH.

**Fig. 5.**
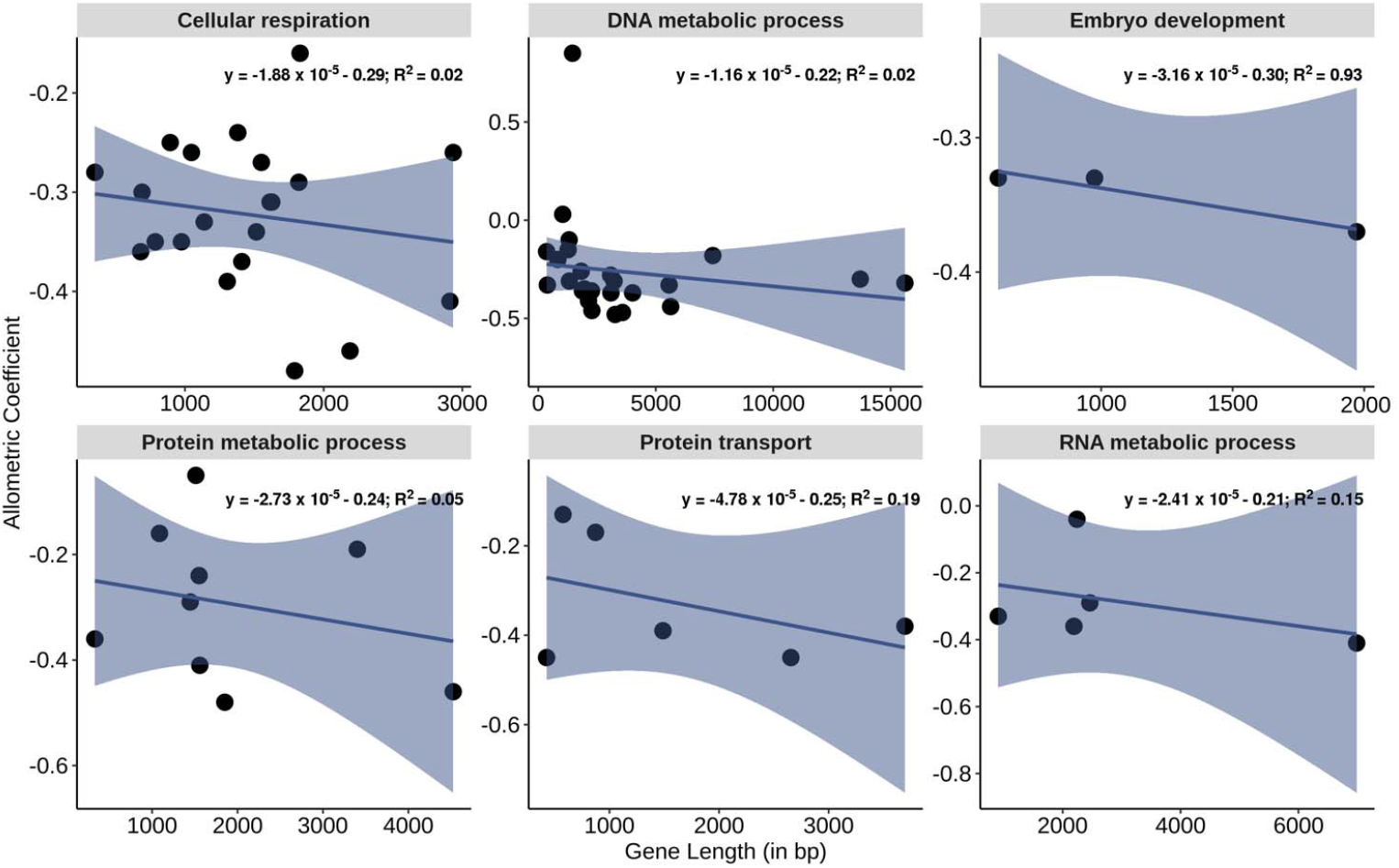
Effect of gene length (in bp) on the allometric coefficient for the scaling of temperature-corrected transcript abundance (in read numbers/g)] and individual body mass (in g) of species from the constructed mock communities.

## Discussion

In this study, we validated the assumptions of the MTE and GRH on the allometric scaling of mRNA transcript and ribosome abundance using constructed mock communities. Using mock communities has been an efficient tool for molecular-based laboratory and computational experiments, leading to thorough quantitative and qualitative validation of new methods (10).

Here, we successfully showed the effect of size and temperature scaling in improving predictive models for the relationship between transcript abundance and biomass to provide a concrete framework in extending the use of metatranscriptomics as a reliable tool in characterizing metazoan communities.

At the genomic level, we found that body size imposes substantial constraints on RNA (mRNA and rRNA) abundance (Fig. 2A), which supports the combined assumptions of GRH and MTE. This is congruent with the previously observed scaling pattern between whole-body RNA concentration and individual body mass. As once put forward by Gillooly *et al*. (8), the size dependence of whole-body RNA concentration is due to the effect of body size on mitochondrial density and ATP production that is needed for RNA transcription and protein synthesis. This may also explain the steeper slope observed in nuclear (mRNA and ribosome) transcripts that play an essential role in the growth and development of individuals. In contrast, the observed slope (closer to the MTE prediction) for mitochondrial transcripts could be due to its limited role in cellular respiration and is, therefore, a good proxy for metabolic rate (11). Overall, this provides the rationale for using RNA-based markers, like the ribosomal RNA ratio, as a proxy for the growth rate in organisms (13). At the gene level, a weaker size dependence on transcript abundance was noted among mitochondrial and nuclear-encoded genes than observed at the genomic level (Fig. 3). Varying allometric coefficients for each gene contradict the widely accepted MTE assumption that mass-specific metabolic processes scale with body mass to the −0.25 power for an interspecific data set (14). The average allometric coefficient for the orthologous genes grouped according to their respective GO terms showed similar patterns. This study shows that genomic and gene-level allometric coefficients may range from −0.30 to −1, which contradicts the constant universal scaling proposed by MTE. This scaling slope (−⅓ to −1) has also been reported for of mass-specific metabolic rate in different organisms under various environmental conditions (15–16).

Furthermore, the correlation between temperature and mass-corrected transcript abundance contradicts the MTE prediction (Fig. 2B). It has been observed that varying factors for transcription of each gene make different genes respond differently to temperature fluctuations (17). The decrease in transcript abundance at low (10 °C) and high (28 °C) incubation temperatures (Fig. S2) demonstrates temperature adaptation of organisms that emphasizes the fundamental thermal kinetics constraints beyond the optimal temperature for growth (18). In terms of the effect of DNA content or genome size in the transcription process, the cell size and nuclear volume may play a critical role. The increase in cell size subsequently requires increased nuclear volume to allow enough nuclear pores through which RNA can pass into the cytoplasm. Most often, selection on the organism’s genome size directs changes in cell size, where an increase in genome size (DNA content) leads to increasing nuclear volume. The genetic basis of GRH (7) suggests that genome size, mainly gene length, appears to be another variable regulating RNA abundance in organisms (Figs. 4 and 5). High copy numbers of internal transcribed spacers and intergenic spacers in genes play structural and catalytic functions during nucleic acid metabolism, carrying elements necessary for mRNA transcription and ribosome production (19). Differences in the number of these repeating units within the genome have been found to have a high correlation with a high RNA concentration (8). The increase in noncoding sequences is most often responsible for the C-value (genome size) increase; thus, interspecific allometries of metabolic processes, e.g., mRNA transcription and ribosome production, can be by-products of evolutionary diversification on the genome size of organisms.

In summary, strong size (biomass) dependence was observed on interspecific scaling of RNA abundance within the constructed mock community. Genome size may also indirectly affect the amount of RNA in tissue by affecting cell volume and overall body size (20). If proven to be valid, these assumptions provide an explanation that incorporates selection pressures acting simultaneously on different levels of biological organization, from genome size, nuclear volume, and cell size up to whole-body mass. This study also highlights the possible use of allometric scaling via a power function as a framework for modeling mRNA transcript and ribosome abundance. We have demonstrated that using allometric scaling coefficients as the power of individual body mass substantially improves the relationship between RNA abundance and species biomass (3). This study provides a robust empirical framework explaining the relationship between RNA abundance through metatranscriptomic reads and the size distribution of species in community samples. However, this needs further validation using actual field samples to fully extend the application of metatranscriptomics for estimating species abundance and active biological processes for community ecology studies.

## Materials and Methods

### Incubation of organisms

A general summary of the entire methodological flow for this study is shown in Fig. S1. Five model species were independently incubated at three temperatures: 10, 19, and 28 °C. For *Caenorhabditis elegans* (ca 1 μg), *Daphnia magna* (ca 5 μg), *Drosophila melanogaster* (ca 10,000 μg), adult individuals were acclimatized at the given temperatures for seven days. After acclimatization, *C. elegans* and *D. melanogaster* eggs were isolated and continuously reared until individuals reached the early adult stage. For *D. magna*, neonate individuals were picked and allowed to grow until the early adult stage. Early adult *Solenopsis invicta* (ca 1,000 μg) workers (ensemble colony) and *Danio rerio* (ca 40,000 μg) were sorted for acclimatization at the same temperatures for seven days. After the acclimatization period, healthy individuals were sorted and continuously reared for one week. In this study, only the early adult individuals were picked for RNA extraction to avoid the potential variations due to the presence of eggs or developing embryos. The number and total wet weight of individuals used per species are shown in Table S1. Three biological replicates incubated at each temperature were prepared in this study.

### Mock community RNA extraction

Collected individuals of each species were separately homogenized in TriPure (Roche, Switzerland) isolation reagent (1 mL/100 mg of the sample) using a tissue homogenizer. The homogenized tissues of each species were then mixed thoroughly to construct a pool of mock community total RNA materials. Next, chloroform (200 μL/mL of TriPure reagent) was added to the pooled solution of homogenized tissues and shaken vigorously for 15 s. The sample was set aside for 30 min at room temperature and then spun in a centrifuge (12,000 *g* for 15 min at 4 °C) to separate the solution into two phases. A total of 600 μL of the resulting upper aqueous phase was transferred to a new tube containing an equal volume of 99.8% ethanol. The sample was then run through the PureLink Mini Kit (Invitrogen, USA) column and spun in the centrifuge at 12,000 *g* for 1 min at room temperature. The column was transferred to a new collection tube and washed twice with 500 μL of Buffer II from the kit. The column membrane was then dried through centrifugation at 12,000 *g* for 1 min in the room. Lastly, the column was transferred to a new recovery tube, and 100 μL of RNAse-free water was added to elute the RNA from the membrane. The quality and concentration of all extracted total RNA samples were analyzed using Bioanalyzer RNA 6000 nano (Agilent Technologies, USA) to measure RNA integrity number (RIN: 21). All samples were stored at −80 °C until the next part of the procedure.

### Library preparation and sequencing

Two types of libraries were prepared from the extracted total RNA of the mock community sample: (1) total RNA library that mainly consists of rRNA and (2) mRNA library isolated from the total RNA sample through poly(A) tail isolation. For total RNA libraries, 1000 ng of extracted total RNA was treated with TURBO^™^ Dnase (ThermoFisher Scientific, USA) to remove traces of DNA materials before cDNA synthesis. Meanwhile, 1000 ng of extracted total RNA was used to prepare mRNA libraries using NEBNext Poly(A) mRNA magnetic isolation (E7490) module (New England BioLabs Inc., USA). Sequencing libraries for total RNA and mRNA were prepared using NEBNext Ultra II RNA Library Prep Kit for Illumina (E7770) and NEBNext Multiplex Oligos (96 Unique Dual Index Primer Pairs) for Illumina following the manufacturer’s protocol. Both total RNA and mRNA were fragmented to *ca* 500-bp lengths, and final enrichment was performed for 8 cycles. The enriched products were then purified using 0.9X Agencourt AMPure XP. Lastly, an equal concentration of the purified libraries was pooled together and sent for Illumina NextSeq 2000 100 SE sequencing (1% PhiX spike-in and 10 pM loading concentration for all libraries) at the NGS High Throughput Genomics Core at the Biodiversity Research Center, Academia Sinica, Taiwan.

### Genomic-level mRNA transcript and ribosome abundance quantification

Quality filtering and adapter removal of both mRNA and total RNA libraries were done using Cutadapt (ver. 2.10, 22). Removal of existing rRNA sequence contaminants in the mRNA libraries was carried out by aligning quality-filtered reads to the reference noncoding RNA sequence of each species (ensemble.org, Table S2) using the BBDuk function within BBTools (https://sourceforge.net/projects/bbmap/). The removal of existing mRNA sequence contaminants in the total RNA libraries was carried out by aligning quality-filtered reads to the reference cDNA consisting of transcript sequences for each species (ensemble.org, Table S2). The remaining reads (Table S3) for each library were normalized by equalizing the number of reads using Seqtk (https://github.com/lh3/seqtk).

To determine the mRNA transcript and ribosome (deduced from rRNA transcripts) abundance for the mitochondrial and nuclear genomes, normalized mRNA and total RNA library reads were aligned to each species’ annotated reference genome sequence using STAR RNA-Seq aligner (ver. 2.7.6, 23). To avoid the mitochondrial transcript being aligned onto a nuclear-encoded mitochondrial pseudogene sequence, mitochondrial genome transcript abundance was determined first through the alignment of library reads to their reference mitochondrial genome (ensemble.org, Table S2). Next, unused reads from this step were placed in a separate fasta file and aligned to each species’ annotated nuclear genome sequence to count the transcript abundance for the genome. Using the indexed BAM file from the read alignment, the number of aligned transcript reads for both the mitochondrial and nuclear genomes were determined by counting the number of aligned reads to the reference sequence for each species using Samtools (24).

Mass-specific transcript read abundance (R) was calculated by dividing the reads counts by the individual body mass per species. From this, temperature-corrected transcript abundance (Re^*E*/*kT*^) was calculated. On the other hand, mass-corrected transcript abundance was calculated as R*M*^α^, where α is the allometric coefficient predicted to be −0.25 according to the MTE (11). Metabolic scaling between ln-transformed temperature-corrected transcript abundance vs individual body mass, ln-transformed temperature-corrected transcript abundance vs genome size (as DNA content collected from https://genomesize.com/) (25), and ln-transformed mass-corrected transcript abundance vs inverse of temperature (1/*kT*) was validated using the lm() function in the R platform (26). Lastly, an ANCOVA was performed to determine the effects of temperature, body size (in g), the genome size (as DNA content), and their interactions on transcript read abundance, using transcript type (mitochondrial and nuclear transcripts) as the categorical factor, while temperature, body size, and genome size served as the covariates.

### Gene-level mRNA transcript and ribosome abundance quantification

Aligned reads to 10 mitochondrial protein-coding [*ATP synthase subunit 8* (ATP8), *NADH dehydrogenase subunit 4L* (NADH4L), and *NADH dehydrogenase subunit 6* (NADH6) were excluded due to the low transcript read counts] and two rRNA (*16S* and *12S*) genes were analyzed in this study. The transcript abundance of the nuclear-encoded rRNA component of the ribosome: (1) large (*28S* for *D. magna*, *D. melanogaster*, *S. invicta*, and *D. rerio* and *26S* for *C. elegans*) and (2) small subunit (*18S*) were also measured. The read counts for all protein-coding genes were determined based on the mRNA library alignment, while total RNA libraries were used for rRNA gene read counts. The transcript read abundance per gene was quantified in transcripts per million (TPM) to normalize values by gene length using Stringtie (ver. 2.14, 27).

Aside from these genes, the transcript read abundance of nuclear-encoded orthologous genes for the five species was assessed in this study. The reference protein sequence (ensembl.org, Table S2) for each species was used to identify single-copy orthologous genes using OrthoFinder (ver. 2.5.4, 28). The protein sequences of all identified orthologous genes for each species were separately consolidated in fasta files and translated back into nucleotide sequences using the EMBOSS Backtranseq tool (ver. 1.12, 29). The resulting nucleotide sequences were then indexed using the bowtie2-build command (30) to serve as the reference in mapping back the normalized reads of mRNA libraries. The transcript read abundance for each orthologous gene was quantified in TPM with RSEM (ver. 1.3.3; 31, 30). A total of 1093 single-copy orthologous genes were identified. These genes were further narrowed down to 62 genes with enough aligned reads for the five species to have sufficient data points for the analysis. Lastly, the functional annotation of the screened orthologous gene was carried out with the EggNOG Mapper (http://eggnog-mapper.embl.de) and DAVID Bioinformatics database (david.abcc.ncifcrf.gov). The allometric coefficient (α) for each gene was calculated by determining the slope from the linear regression between temperature-corrected transcript abundance for each gene vs individual body mass using the lm() function in R.

To see if transcript scaling can improve predictive models for the relationship between RNA transcripts and biomass, we ran several sets of correlation analyses among commonly used gene markers [nuclear (*18S* and *28S*) and mitochondrial (*12S*, *16S*, Cytochrome oxidase subunit I, and Cytochrome b apoenzyme) markers] for community ecology studies. Comparison between the adjusted *R* (*R*^2^) among three different analyses was made in this study: (1) ln-relationship between uncorrected mass-specific transcript read number [(R) (reads/g)] and individual body mass (in g), which excludes the effect of temperature and body size in scaling; (2) ln-relationship between temperature-corrected transcript abundance ([Re^*E*/*kT*^] (in read numbers/g)) and individual body mass (in g) to see the effect of temperature scaling (32); and (3) ln-relationship between mass-specific transcript read number (reads/g) and allometrically scaled biomass (*M*^α^ [in g]), where α refers to the identified allometric coefficient for each gene, (Table S4) for body size scaling (3).

## Acknowledgments

The authors thank the Taiwan Zebrafish Core Facility at Academia Sinica (TZCAS) at the Institute of Cellular and Organismic Biology and NGS High Throughput Genomics Core at the Biodiversity Research Center, Academia Sinica. Lastly, we would like to acknowledge all helpful feedback and comments from Dr. Jason I-sheng Tsai, Dr. John Wang, Dr. Michael Monaghan, and Dr. Chih-Hao Hsieh.

## Funding

This study was supported by the following grants:

Taiwan International Graduate Program, Academia Sinica and National Taiwan Normal University (MLDL)

Academia Sinica, Taiwan (RJM)

Ministry of Science and Technology, Taiwan 108-2611-M-001 and 109-2611-M491 001 (RJM)

Scientific Committee on Oceanic Research working group 157 (RJM)

## Author contributions

The following are the authors’ contributions:

Conceptualization: MLDL, RJM, SQS

Methodology: MLDL, RJM, YYL

Investigation: MLDL, RJM

Visualization: MLDL, RJM

Supervision: RJM

Writing—original draft: MLDL, RJM

Writing—review & editing: MLDL, RJM, SQS

## Data and materials availability

The raw sequences will be deposited in DNA Data Bank of Japan (DDBJ). Currently, DDBJ submission services are under maintenance and will be running upon further notice. Codes used in data analyses can be accessed thru https://github.com/mldlopez/Allometric-scaling-of-RNA-abundance-from-genes-to-communities/blob/master/codes.pdf. All data are available in the main text and/or the supplementary materials.

## References

1. R. W. Sterner, J. J. Elser. Ecological Stoichiometry: The Biology of Elements from Molecules to the Biosphere (Princeton University Press, Princeton, NJ, 2002).

2. R. M. Sibly, J. H. Brown, A. Kodric-Brown. Metabolic Ecology: a Scaling Approach (Wiley-Blackwell, Oxford, 2012).

3. M. C. Yates, D. Glaser, J. Post, M. E. Cristescu, D. J. Fraser, A. M. Derry. The relationship between eDNA particle concentration and organism abundance in nature is strengthened by allometric scaling. Mol. Ecol. 30, 3068–3082 (2020).

4. J. H. Brown, J. F. Gillooly, A. P. Allen, V. M. Savage, G. B. West. Toward a metabolic theory of ecology. Ecol., 85, 1771–1789 (2004).

5. J. J. Elser, R. W. Sterner, E. Gorokhova, W. F. Fagan, T. A. Markow, J. B. Cotner, J. Harrison, S. E. Hobbie, G. M. Odell, J. W. Weider. Biological stoichiometry from genes to ecosystems. Ecol. Lett., 3, 540–550 (2008).

6. F. J. Bullejos, P. Carrillo, E. Gorokhova, J. M. Medina-Sánchez, M. Villar-Argaiz. Nucleic acid content in crustacean zooplankton: bridging metabolic and stoichiometric predictions. PLoS ONE, 9(1), e86493 (2014).

7. G. Grimaldi, P. O. Di Nocera. Multiple repeated units in *Drosophila melanogaster* ribosomal DNA spacer stimulate rRNA precursor transcription. Proc. Natl. Acad. Sci. U.S.A., 85, 5502–5506 (1988).

8. J. F. Gillooly, A. P. Allen, J. H. Brown, J. J. Elser, C. M. del Rio, V. M. Savage, G. B. West, W. H. Woodruff, H. A. Woods. The metabolic basis of whole-organism RNA and phosphorus stoichiometry, Proc. Natl. Acad. Sci. U.S.A., 102, 11923–11927 (2005).

9. A. P. Allen, J. F. Gillooly. Towards integration of ecological stoichiometry and the metabolic theory of ecology to better understand nutrient cycling. Ecol. Lett., 12, 369–384 (2009).

10. M. L. D. Lopez, Y. Y. Lin, M. Sato, C. H. Hsieh, F. K. Shiah, R. J. Machida. Using metatranscriptomics to estimate the diversity and composition of zooplankton communities. Mol. Ecol. Resour. doi: 10.1111/1755-0998.13506.

11. J. F. Gillooly, J. H. Brown, G. B. West, V. M. Savage, E. L. Charnov. Effects of size and temperature on metabolic rate. Science, 293, 2248–2251 (2001).

12. A. Ben-Shem, N. G. de Loubresse, S. Melnikov, L. Jenner, G. Yusupova, M. Yusupov. The structure of the eukaryotic ribosome at 3.0 Å resolution. Science, 334, 1524–1529 (2011).

13. W. L. Kong, T. Miki, Y. Y. Lin, W. Makino, J. Urabe, S. H. Gu, R.J. Machida. Nuclear and mitochondrial ribosomal ratio as an index of animal growth rate. Limnol. Oceanogr. Methods, 17(11), 575–584 (2019).

14. R. H. Peters. The Ecological Implications of Body Size (Cambridge University Press, Cambridge, 1983).

15. D. S. Glazier. Metabolic scaling in complex systems. Systems, 2, 451–540 (2014).

16. C.R. White, M. R. Kearney. Metabolic scaling in animals: Methods, empirical results, and theoretical explanations. Compr. Physiol., 4, 231–256 (2014).

17. S. M. D. Oliveira, A. Häkkinen, J. Lloyd-Price, H. Tran, V. Kandavalli, A. S. Ribeiro. Temperature-dependent model of multi-step transcription initiation in *Escherichia coli* based on live single-cell measurements. PLoS Comput. Biol., 12(10), e1005174 (2016).

18. A. Clarke. Costs and consequences of evolutionary temperature adaptation. Trends in Ecology and Evolution, 18, 573–581 (2003).

19. R. H. Reeder, M. Dunaway. Spacer regulation of xenopus ribosomal gene transcription. Competition in oocytes. Cell, 35, 449–456 (1983).

20. J. Kozłowski, M. Konarzewski, A. T. Gawelczyk. Cell size as a link between non-coding DNA and metabolic rate scaling. Proc. Natl. Acad. Sci. U.S.A., 100(24), 14080–14085 (2003).

21. A. Schroeder, O. Mueller, S. Stocker, R. Salowsky, M. Leiber, M. Gassmann, S. Lightfoot, W. Menzel, M. Granzow, T. Ragg. The RIN: An RNA integrity number for assigning integrity values to RNA measurements. BMC Mol. Biol., 7, 3 (2006).

22. M. Martin. Cutadapt removes adapter sequences from high-throughput sequencing reads. EMBnet.Journal, 17(1), 10–12 (2011).

23. A. Dobin, C. A. Davis, F. Schlesinger, J. Drenkow, C. Zaleski, S. Jha, P. Batut, M. Chaisson, T. R. Gingeras. STAR: ultrafast universal RNA-seq aligner. Bioinf., 29(1), 15–21 (2013).

24. H. Li, B. Handsaker, A. Wysoker, T. Fennell, J. Ruan, N. Homer, G. Marth, G. Abecasis, R. Durbin. The Sequence Alignment/Map format and SAMtools, Bioinf., 25(16), 2078–2079 (2009).

25. T. R. Gregory, J. A. Nicol, H. Tamm, B. Kullman, K. Kullman, J. J. Leitch, B. G. Murray, D. F. Kapraun, J. Greilhuber, M. D. Bennett. Eukaryotic genome size databases, Nucleic Acids Re., 35(1), 332–338 (2007).

26. R Core Team R: A Language and Environment for Statistical Computing, Vienna, Austria. Available at: https://www.R-project.org/ (2017).

27. M. Pertea, G. Pertea, C. Antonescu, T. C. Chang, J. T. Mendell, S. L. Salzberg. StringTie enables improved reconstruction of a transcriptome from RNA-seq reads. Nat. Biotech., 33, 290–295 (2015).

28. D. M. Emms, S. Kelly. OrthoFinder: phylogenetic orthology inference for comparative genomics. Genome Biol., 20, 238 (2019).

29. F. Madeira, Y. M. Park, J. Lee, N. Buso, T. Gur, N. Madhusoodanan, P. Basutkar, A. R. N. Tivey, P. C. Potter, R. D. Finn, R. Lopez. The EMBL-EBI search and sequence analysis tools APIs in 2019. Nucleic Acids Res., 47(W1), W636–W641 (2019).

30. B. Langmead, S. Salzberg. Fast gapped-read alignment with Bowtie 2. Nat. Methods, 9, 357–359 (2012).

31. B. Li, C. N. Dewey. RSEM: Accurate transcript quantification from RNA-Seq data with or without a reference genome. BMC Bioinf., 12, 323 (2011).

32. J. F. Gillooly, A. P. Allen, G. B. West, J. H. Brown. The rate of DNA evolution: Effects of body size and temperature on the molecular clock. Proc. Natl. Acad. Sci. U.S.A., 102(1), 140–145 (2005).

