## Supplemental Materials for "Allometric scaling of RNA abundance from genes to communities"

**This PDF file includes:**

Figs. S1 and S2

Tables S1 to S5

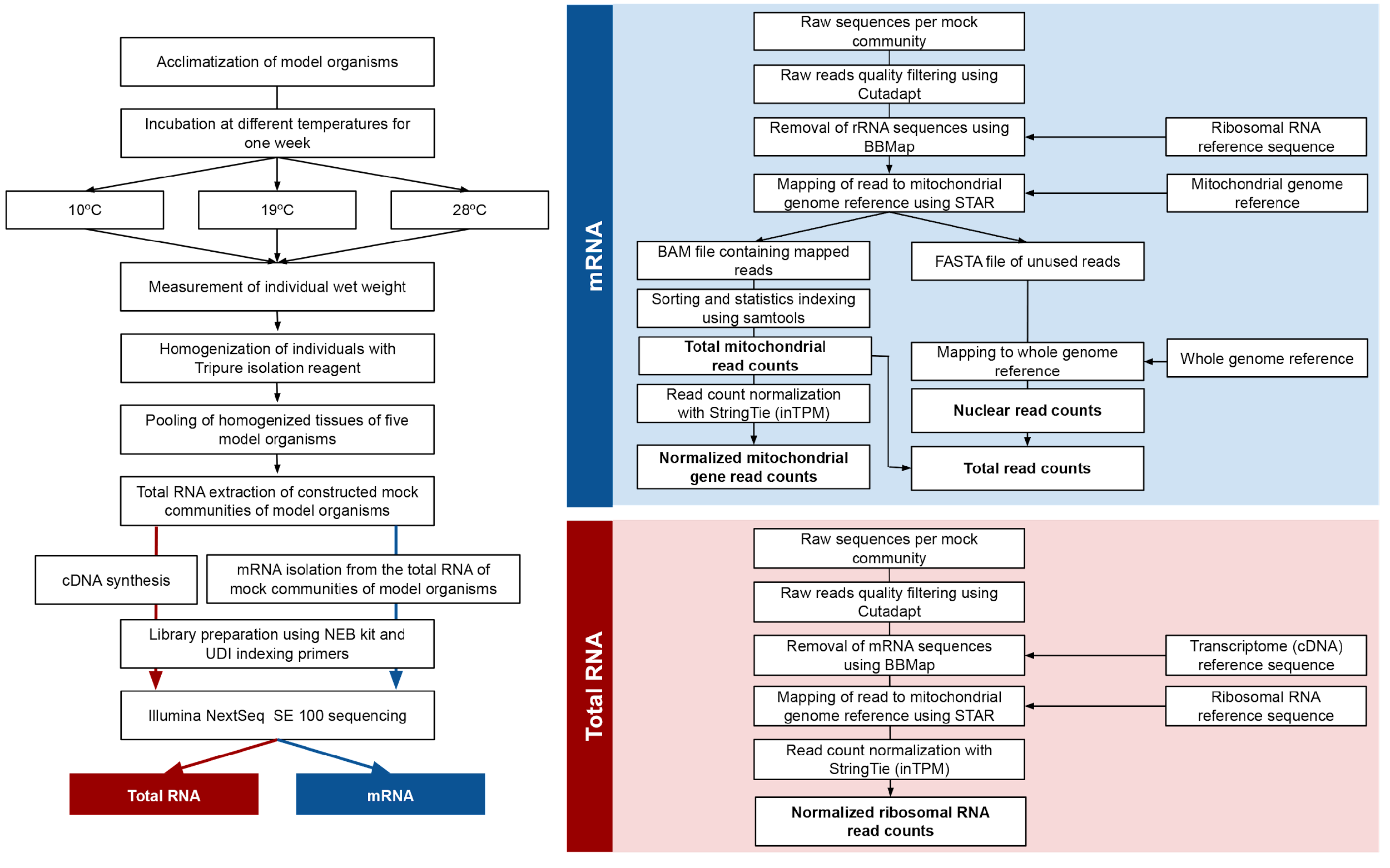

**Fig. S1.** General summary for the methodological workflow of the study. Bold text shows the measured reads from the mRNA and total RNA library alignment.

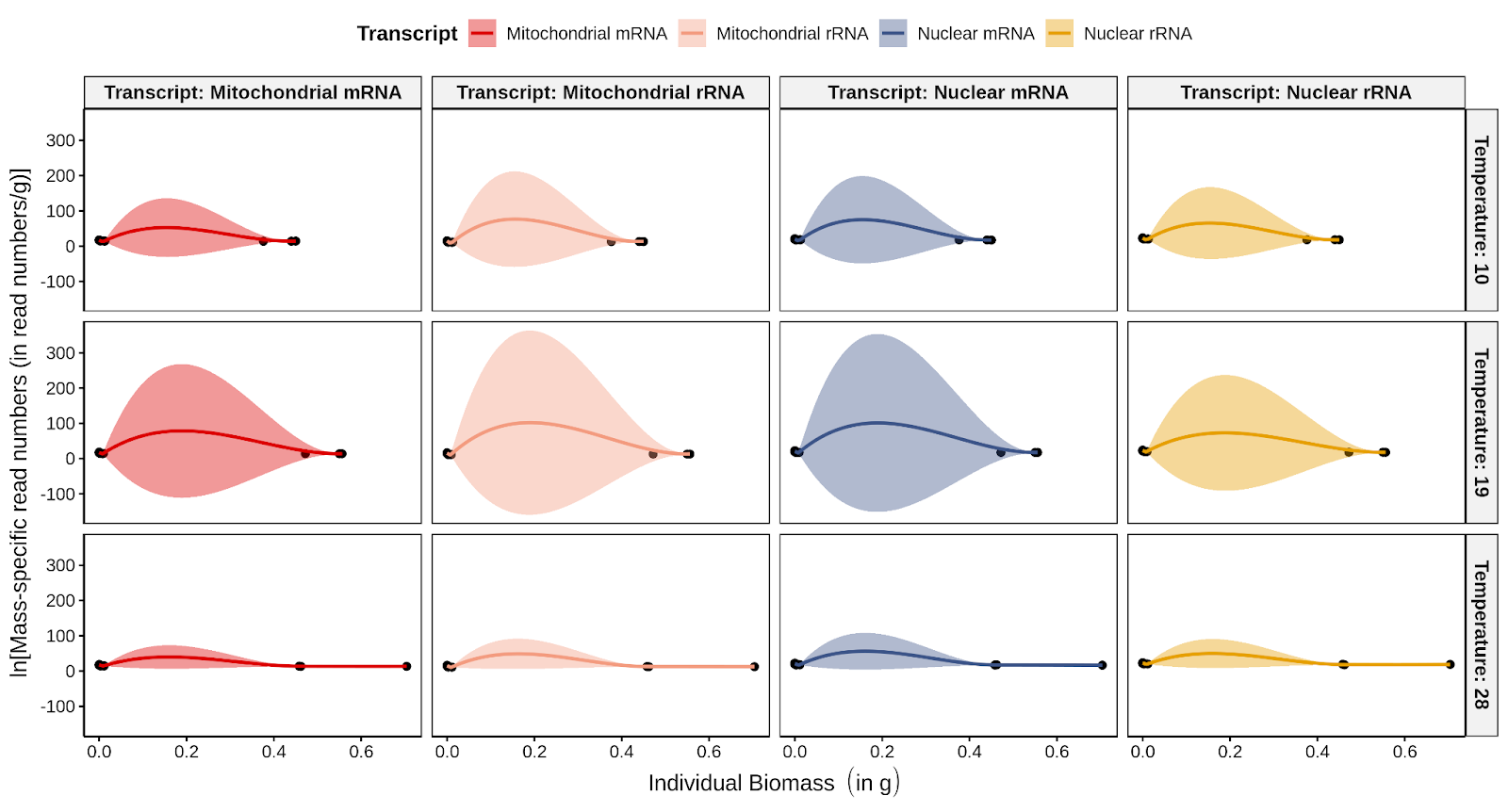

**Fig. S2.** Scatter plot for the effect of temperature (°C) and biomass (in g) on temperature-corrected transcript abundance [[Re^(^*^E^*^/^*^kT^*^)^] (in read numbers/g)] of species.

**Table S1.** Summary for the number of individual and total wet weight (in g) used in constructing the mock communities.

| Trial | Temperature  (in °C) | Species | Number of individuals | Total biomass (in g) |
| --- | --- | --- | --- | --- |
| 1 | 10 | *Caenorhabditis elegans* | 500 | 0.0005 |
|  |  | *Daphnia magna* | 30 | 0.00045 |
|  |  | *Solenopsis invicta* | 15 | 0.043 |
|  |  | *Drosophila melanogaster* | 10 | 0.114 |
|  |  | *Danio rerio* | 1 | 0.45 |
|  | 19 | *Caenorhabditis elegans* | 500 | 0.0005 |
|  |  | *Daphnia magna* | 30 | 0.00039 |
|  |  | *Solenopsis invicta* | 15 | 0.037 |
|  |  | *Drosophila melanogaster* | 10 | 0.078 |
|  |  | *Danio rerio* | 1 | 0.55 |
|  | 28 | *Caenorhabditis elegans* | 500 | 0.0005 |
|  |  | *Daphnia magna* | 30 | 0.00042 |
|  |  | *Solenopsis invicta* | 15 | 0.035 |
|  |  | *Drosophila melanogaster* | 10 | 0.066 |
|  |  | *Danio rerio* | 1 | 0.458 |
| 2 | 10 | *Caenorhabditis elegans* | 500 | 0.0005 |
|  |  | *Daphnia magna* | 30 | 0.00033 |
|  |  | *Solenopsis invicta* | 15 | 0.035 |
|  |  | *Drosophila melanogaster* | 10 | 0.118 |
|  |  | *Danio rerio* | 1 | 0.44 |
|  | 19 | *Caenorhabditis elegans* | 500 | 0.0005 |
|  |  | *Daphnia magna* | 30 | 0.00036 |
|  |  | *Solenopsis invicta* | 15 | 0.041 |
|  |  | *Drosophila melanogaster* | 10 | 0.098 |
|  |  | *Danio rerio* | 1 | 0.472 |
|  | 28 | *Caenorhabditis elegans* | 500 | 0.0005 |
|  |  | *Daphnia magna* | 30 | 0.00039 |
|  |  | *Solenopsis invicta* | 15 | 0.043 |
|  |  | *Drosophila melanogaster* | 10 | 0.112 |
|  |  | *Danio rerio* | 1 | 0.704 |
| 3 | 10 | *Caenorhabditis elegans* | 500 | 0.0005 |
|  |  | *Daphnia magna* | 30 | 0.00042 |
|  |  | *Solenopsis invicta* | 15 | 0.039 |
|  |  | *Drosophila melanogaster* | 10 | 0.13 |
|  |  | *Danio rerio* | 1 | 0.376 |
|  | 19 | *Caenorhabditis elegans* | 500 | 4 |
|  |  | *Daphnia magna* | 30 | 0.00045 |
|  |  | *Solenopsis invicta* | 15 | 0.036 |
|  |  | *Drosophila melanogaster* | 10 | 0.082 |
|  |  | *Danio rerio* | 1 | 0.556 |
|  | 28 | *Caenorhabditis elegans* | 500 | 0.0005 |
|  |  | *Daphnia magna* | 30 | 0.00045 |
|  |  | *Solenopsis invicta* | 15 | 0.044 |
|  |  | *Drosophila melanogaster* | 10 | 0.112 |
|  |  | *Danio rerio* | 1 | 0.462 |

**Table S2.** FTP links of the reference sequences for the model species used in the study.

| **Species** | **Reference sequence** | **FTP link** |
| --- | --- | --- |
| *Caenorhabditis elegans* | *Genomic* | [*http://ftp.ebi.ac.uk/ensemblgenomes/pub/release-51/metazoa/fasta/caenorhabditis_elegans/dna/*](http://ftp.ebi.ac.uk/ensemblgenomes/pub/release-51/metazoa/fasta/caenorhabditis_elegans/dna/) |
|  | *cDNA* | [*http://ftp.ebi.ac.uk/ensemblgenomes/pub/release-51/metazoa/fasta/caenorhabditis_elegans/cdna/*](http://ftp.ebi.ac.uk/ensemblgenomes/pub/release-51/metazoa/fasta/caenorhabditis_elegans/cdna/) |
|  | *Non-coding RNA* | [*http://ftp.ebi.ac.uk/ensemblgenomes/pub/release-51/metazoa/fasta/caenorhabditis_elegans/ncrna/*](http://ftp.ebi.ac.uk/ensemblgenomes/pub/release-51/metazoa/fasta/caenorhabditis_elegans/ncrna/) |
|  | *Protein* | [*http://ftp.ebi.ac.uk/ensemblgenomes/pub/release-51/metazoa/fasta/caenorhabditis_elegans/pep/*](http://ftp.ebi.ac.uk/ensemblgenomes/pub/release-51/metazoa/fasta/caenorhabditis_elegans/pep/) |
| *Daphnia magna* | *Genomic* | [*http://ftp.ebi.ac.uk/ensemblgenomes/pub/release-51/metazoa/fasta/daphnia_magna/dna/*](http://ftp.ebi.ac.uk/ensemblgenomes/pub/release-51/metazoa/fasta/daphnia_magna/dna/) |
|  | *cDNA* | [*http://ftp.ebi.ac.uk/ensemblgenomes/pub/release-51/metazoa/fasta/daphnia_magna/cdna/*](http://ftp.ebi.ac.uk/ensemblgenomes/pub/release-51/metazoa/fasta/daphnia_magna/cdna/) |
|  | *Non-coding RNA* | [*http://ftp.ebi.ac.uk/ensemblgenomes/pub/release-51/metazoa/fasta/daphnia_magna/ncrna/*](http://ftp.ebi.ac.uk/ensemblgenomes/pub/release-51/metazoa/fasta/daphnia_magna/ncrna/) |
|  | *Protein* | [*http://ftp.ebi.ac.uk/ensemblgenomes/pub/release-51/metazoa/fasta/daphnia_magna/pep/*](http://ftp.ebi.ac.uk/ensemblgenomes/pub/release-51/metazoa/fasta/daphnia_magna/pep/) |
| *Solenopsis invicta* | *Genomic* | [*http://ftp.ebi.ac.uk/ensemblgenomes/pub/release-51/metazoa/fasta/solenopsis_invicta/dna/*](http://ftp.ebi.ac.uk/ensemblgenomes/pub/release-51/metazoa/fasta/solenopsis_invicta/dna/) |
|  | *cDNA* | [*http://ftp.ebi.ac.uk/ensemblgenomes/pub/release-51/metazoa/fasta/solenopsis_invicta/cdna/*](http://ftp.ebi.ac.uk/ensemblgenomes/pub/release-51/metazoa/fasta/solenopsis_invicta/cdna/) |
|  | *Non-coding RNA* | [*http://ftp.ebi.ac.uk/ensemblgenomes/pub/release-51/metazoa/fasta/solenopsis_invicta/ncrna/*](http://ftp.ebi.ac.uk/ensemblgenomes/pub/release-51/metazoa/fasta/solenopsis_invicta/ncrna/) |
|  | *Protein* | [*http://ftp.ebi.ac.uk/ensemblgenomes/pub/release-51/metazoa/fasta/solenopsis_invicta/pep/*](http://ftp.ebi.ac.uk/ensemblgenomes/pub/release-51/metazoa/fasta/solenopsis_invicta/pep/) |
| *Drosophila melanogaster* | *Genomic* | [*http://ftp.ebi.ac.uk/ensemblgenomes/pub/release-51/metazoa/fasta/drosophila_melanogaster/dna/*](http://ftp.ebi.ac.uk/ensemblgenomes/pub/release-51/metazoa/fasta/drosophila_melanogaster/dna/) |
|  | *cDNA* | [*http://ftp.ebi.ac.uk/ensemblgenomes/pub/release-51/metazoa/fasta/drosophila_melanogaster/cdna/*](http://ftp.ebi.ac.uk/ensemblgenomes/pub/release-51/metazoa/fasta/drosophila_melanogaster/cdna/) |
|  | *Non-coding RNA* | [*http://ftp.ebi.ac.uk/ensemblgenomes/pub/release-51/metazoa/fasta/drosophila_melanogaster/ncrna/*](http://ftp.ebi.ac.uk/ensemblgenomes/pub/release-51/metazoa/fasta/drosophila_melanogaster/ncrna/) |
|  | *Protein* | [*http://ftp.ebi.ac.uk/ensemblgenomes/pub/release-51/metazoa/fasta/drosophila_melanogaster/pep/*](http://ftp.ebi.ac.uk/ensemblgenomes/pub/release-51/metazoa/fasta/drosophila_melanogaster/pep/) |
| *Danio rerio* | *Genomic* | [*http://ftp.ensembl.org/pub/release-104/fasta/danio_rerio/dna/*](http://ftp.ensembl.org/pub/release-104/fasta/danio_rerio/dna/) |
|  | *cDNA* | [*http://ftp.ensembl.org/pub/release-104/fasta/danio_rerio/cdna/*](http://ftp.ensembl.org/pub/release-104/fasta/danio_rerio/cdna/) |
|  | *Non-coding RNA* | [*http://ftp.ensembl.org/pub/release-104/fasta/danio_rerio/ncrna/*](http://ftp.ensembl.org/pub/release-104/fasta/danio_rerio/ncrna/) |
|  | *Protein* | [*http://ftp.ensembl.org/pub/release-104/fasta/danio_rerio/pep/*](http://ftp.ensembl.org/pub/release-104/fasta/danio_rerio/pep/) |

**Table S3.** Summary for the number of individuals and total wet weight (in g) used in constructing the mock communities.

| Trial | Temperature  (in °C) | Library type | Raw count | Quality filtering | Sequence contaminant removal | Read number normalization |
| --- | --- | --- | --- | --- | --- | --- |
| 1 | 10 | mRNA | 28,678,097 | 28,678,097 | 27,297,999 | 24,765,099 |
|  | 19 |  | 28,678,097 | 28,678,097 | 24,906,652 | 24,765,099 |
|  | 28 |  | 25,895,726 | 25,895,726 | 25,036,498 | 24,765,099 |
| 2 | 10 |  | 29,049,006 | 29,049,006 | 28,112,868 | 24,765,099 |
|  | 19 |  | 31,616,488 | 31,616,488 | 30,365,611 | 24,765,099 |
|  | 28 |  | 31,118,045 | 31,118,045 | 29,979,772 | 24,765,099 |
| 3 | 10 |  | 25,788,436 | 25,788,436 | 24,765,099 | 24,765,099 |
|  | 19 |  | 31,413,509 | 31,413,509 | 24,765,099 | 24,765,099 |
|  | 28 |  | 31,413,509 | 31,413,509 | 30,340,297 | 18,299,638 |
| 1 | 10 | total RNA | 31,892,680 | 31,892,680 | 21,850,634 | 18,299,638 |
|  | 19 |  | 30,998,696 | 30,998,696 | 20,230,186 | 18,299,638 |
|  | 28 |  | 30,018,702 | 30,018,702 | 21,696,707 | 18,299,638 |
| 2 | 10 |  | 36,387,892 | 36,387,892 | 23,717,275 | 18,299,638 |
|  | 19 |  | 31,828,942 | 31,828,942 | 21,411,384 | 18,299,638 |
|  | 28 |  | 28,413,054 | 28,413,054 | 18,354,411 | 18,299,638 |
| 3 | 10 |  | 31,316,778 | 31,316,778 | 22,345,844 | 18,299,638 |
|  | 19 |  | 27,871,203 | 27,871,203 | 18,306,138 | 18,299,638 |
|  | 28 |  | 37,713,476 | 37,713,476 | 26,937,424 | 18,299,638 |

**Table S4.** Parameter estimates for the regressions between the natural logarithm (ln) of temperature-corrected transcript abundance [ln(Re*^E^*^/^*^kT^*) vs individual body mass (in g) of selected mitochondrial and nuclear genes.

| Source | Gene | ln(Re*^E^*^/^*^kT^*) vs ln(*M*) | | | | |
| --- | --- | --- | --- | --- | --- | --- |
|  |  | Fitted slope ± 95% CI | Predicted slope | Intercept | Adjusted *R*^2^ | *P*-value |
| Nuclear ribosomal RNA | 18S | –0.48 ± 0.02 | –0.25 | 37.84 | 0.86 | **<0.01** |
|  | 28S | –0.46 ± 0.04 | –0.25 | 38.30 | 0.69 | **<0.01** |
| Mitochondrial protein coding genes & ribosomal RNA | 12S | –0.41 ± 0.07 | –0.25 | 33.27 | 0.43 | **<0.01** |
|  | 16S | –0.24 ± 0.03 | –0.25 | 34.69 | 0.53 | **<0.01** |
|  | ATP6 | –0.36 ± 0.03 | –0.25 | 36.70 | 0.80 | **<0.01** |
|  | COI | –0.27 ± 0.02 | –0.25 | 37.94 | 0.73 | **<0.01** |
|  | COII | –0.30 ± 0.02 | –0.25 | 37.54 | 0.71 | **<0.01** |
|  | COIII | –0.35 ± 0.02 | –0.25 | 37.45 | 0.85 | **<0.01** |
|  | CytB | –0.33 ± 0.02 | –0.25 | 37.01 | 0.76 | **<0.01** |
|  | NADH1 | –0.35 ± 0.03 | –0.25 | 35.70 | 0.69 | **<0.01** |
|  | NADH2 | –0.26 ± 0.04 | –0.25 | 35.19 | 0.46 | **<0.01** |
|  | NADH3 | –0.28 ± 0.02 | –0.25 | 34.66 | 0.70 | **<0.01** |
|  | NADH4 | –0.24 ± 0.03 | –0.25 | 36.01 | 0.60 | **<0.01** |
|  | NADH5 | –0.29 ± 0.02 | –0.25 | 35.66 | 0.71 | **<0.01** |

**Table S5.** Parameter estimates for the regressions between the natural logarithm (ln) of temperature-corrected transcript abundance [ln(Re*^E^*^/^*^kT^*) vs individual body mass (in g) of identified nuclear-encoded orthologous genes for the five species.

| GO terms | Gene code | Gene name | ln(Re*^E^*^/^*^kT^*) vs ln(*M*) | | | | |
| --- | --- | --- | --- | --- | --- | --- | --- |
|  |  |  | Fitted slope ± 95% CI | Predicted slope | Intercept | Adjusted *R*^2^ | *P*-value |
| Cellular respiration | ABCE1 | ATP-binding cassette, sub-family E (OABP), member 1 | –0.16 ± 0.10 | –0.25 | 34.14 | 0.07 | **>0.01** |
|  | ATP2C1 | ATPase, Ca^++^ transporting, type 2C, member 1 | –0.26 ± 0.60 | –0.25 | 32.77 | 0.07 | **>0.01** |
|  | ATP5A1 | ATP synthase, H^+^ transporting, mitochondrial F1 complex, alpha subunit 1, cardiac muscle | –0.31 ± 0.04 | –0.25 | 38.65 | 0.48 | **<0.01** |
|  | ATP6V1H | ATPase, H^+^ transporting, lysosomal V1 subunit H | –0.37 ± 0.09 | –0.25 | 33.33 | 0.45 | **<0.01** |
|  | CCT5 | Chaperonin containing TCP1, subunit 5 (epsilon) | –0.31 ± 0.07 | –0.25 | 34.79 | 0.35 | **<0.01** |
|  | DDX46 | DEAD (Asp–Glu–Ala–Asp) box polypeptide 46 | –0.41 ± 0.32 | –0.25 | 33.21 | 0.04 | >0.01 |
|  | ETFDH | Electron-transferring-flavoprotein dehydrogenase | –0.48 ± 0.22 | –0.25 | 32.61 | 0.35 | **<0.01** |
|  | MCM7 | Minichromosome maintenance complex component 7 | –0.46 ± 0.12 | –0.25 | 35.51 | 0.28 | **<0.01** |
|  | PSMC2 | Proteasome 26S subunit, ATPase 2 | –0.39 ± 0.07 | –0.25 | 35.73 | 0.36 | **<0.01** |
|  | RTCB | RNA 2′,3′-cyclic phosphate and 5′-OH ligase | –0.34 ± 0.11 | –0.25 | 34.11 | 0.34 | **<0.01** |
|  | SDHB | Succinate dehydrogenase complex, subunit B, iron sulfur (Ip) | –0.25 ± 0.06 | –0.25 | 38.94 | 0.23 | **<0.01** |
| DNA metabolic process | AHCY | Adenosylhomocysteinase | –0.10 ± 0.09 | –0.25 | 38.72 | 0.00 | >0.01 |
|  | AP-50 | AP-50 complex subunit | –0.31 ± 0.13 | –0.25 | 34.61 | 0.14 | **<0.01** |
|  | ARFGAP1 | ADP-ribosylation factor GTPase activating protein 1 | –0.15 ± 0.13 | –0.25 | 36.30 | 0.01 | >0.01 |
|  | CDC5L | CDC5 cell division cycle 5-like (*S. pombe*) | –0.36 ± 0.10 | –0.25 | 33.89 | 0.24 | **<0.01** |
|  | CNOT11 | CCR4-NOT transcription complex, subunit 11 | –0.36 ± 0.16 | –0.25 | 33.58 | 0.22 | **<0.01** |
|  | CPSF3L | Cleavage and polyadenylation specific factor 3-like | –0.26 ± 0.08 | –0.25 | 35.09 | 0.18 | **<0.01** |
|  | DNAJC21 | DnaJ (Hsp40) homolog, subfamily C, member 21 | 0.85 ± 0.36 | –0.25 | 36.48 | 0.36 | **<0.01** |
|  | DNM1L | Dynamin 1-like | –0.41 ± 0.10 | –0.25 | 37.14 | 0.34 | **<0.01** |
|  | DYNC1H1 | Dynein, cytoplasmic 1, heavy chain 1 | –0.30 ± 0.08 | –0.25 | 35.31 | 0.21 | **<0.01** |
|  | EIF3-S10 | Eukaryotic translation initiation factor 3 subunit A | –0.44 ± 0.28 | –0.25 | 31.14 | 0.18 | **<0.01** |
|  | EIF5B | Eukaryotic translation initiation factor 5B | –0.31 ± 0.12 | –0.25 | 34.11 | 0.15 | **<0.01** |
|  | HDC | Histidine decarboxylase | 0.03 ± 0.32 | –0.25 | 34.33 | -0.07 | >0.01 |
|  | HSP90B1 | Heat shock protein 90, beta (grp94), member 1 | –0.46 ± 0.09 | –0.25 | 34.52 | 0.54 | **<0.01** |
|  | PCSK2 | Proprotein convertase subtilisin/kexin type 2 | –0.35 ± 0.11 | –0.25 | 32.74 | 0.33 | **<0.01** |
|  | PNO1 | Partner of NOB1 homolog | –0.20 ± 0.10 | –0.25 | 36.75 | 0.07 | >0.01 |
|  | POLR2A | Polymerase (RNA) II (DNA directed) polypeptide A | –0.33 ± 0.11 | –0.25 | 32.58 | 0.21 | **<0.01** |
|  | POLR2B | Polymerase (RNA) II (DNA directed) polypeptide B | –0.47 ± 0.09 | –0.25 | 32.91 | 0.38 | **<0.01** |
|  | PRKAA2 | Protein kinase, AMP-activated, alpha 2 catalytic subunit | –0.36 ± 0.07 | –0.25 | 34.26 | 0.43 | **<0.01** |
|  | RB1CC1 | RB1-inducible coiled-coil 1 | –0.37 ± 0.46 | –0.25 | 31.12 | 0.05 | >0.01 |
|  | RNF2 | Ring finger protein 2 | –0.18 ± 0.16 | –0.25 | 35.74 | 0.01 | >0.01 |
|  | RPS25 | Ribosomal protein S25 | –0.16 ± 0.10 | –0.25 | 40.27 | 0.04 | >0.01 |
|  | RYR | Ryanodine receptor | –0.32 ± 0.07 | –0.25 | 32.91 | 0.54 | **<0.01** |
|  | SKIV2L2 | Superkiller viralicidic activity 2-like 2 | –0.28 ± 0.34 | –0.25 | 33.82 | 0.02 | >0.01 |
|  | SUPT16H | SPT16 homolog, facilitates chromatin remodeling subunit | –0.37 ± 0.08 | –0.25 | 34.89 | 0.28 | **<0.01** |
|  | UBA52 | Ubiquitin A-52 residue ribosomal protein fusion product 1 | –0.33 ± 0.07 | –0.25 | 40.09 | 0.33 | **<0.01** |
|  | VINC | Vinculin | –0.48 ± 0.07 | –0.25 | 33.09 | 0.68 | **<0.01** |
| Embryo  development | CACTIN | Cactin | –0.37 ± 0.14 | –0.25 | 33.73 | 0.24 | **<0.01** |
|  | CFAP20 | Cilia and flagella associated protein 20 | –0.33 ± 0.10 | –0.25 | 35.79 | 0.22 | **<0.01** |
|  | GNB2I1 | Guanine nucleotide-binding protein (G protein), beta polypeptide 2-like 1 | –0.33 ± 0.09 | –0.25 | 38.79 | 0.24 | **<0.01** |
| Protein metabolic process | ARIH2 | Ariadne homolog 2 | –0.29 ± 0.10 | –0.25 | 35.23 | 0.16 | **<0.01** |
|  | DDB1 | Damage-specific DNA binding protein 1 | –0.19 ± 0.28 | –0.25 | 37.03 | 0.02 | >0.01 |
|  | PSMD3 | Proteasome 26S subunit, non-ATPase 3 | –0.05 ± 0.32 | –0.25 | 35.33 | 0.07 | >0.01 |
|  | PSMD7 | Proteasome 26S subunit, non-ATPase 7 | –0.16 ± 0.10 | –0.25 | 37.40 | 0.04 | >0.01 |
|  | RBX1 | Ring-box 1, E3 ubiquitin protein ligase | –0.36 ± 0.13 | –0.25 | 37.73 | 0.16 | **<0.01** |
| Protein transport | AP2S1 | Adaptor-related protein complex 2, sigma 1 subunit | –0.45 ± 0.15 | –0.25 | 35.30 | 0.29 | **<0.01** |
|  | AP3S2 | Adaptor-related protein complex 3, sigma 2 subunit | –0.13 ± 0.39 | –0.25 | 34.45 | 0.02 | >0.01 |
|  | COPA | Coatomer protein complex, subunit alpha | –0.38 ± 0.10 | –0.25 | 33.04 | 0.33 | **<0.01** |
|  | COPG2 | Coatomer protein complex, subunit gamma 2 | –0.45 ± 0.13 | –0.25 | 33.53 | 0.22 | **<0.01** |
|  | SRP54 | Signal recognition particle 54 | –0.39 ± 0.07 | –0.25 | 34.68 | 0.41 | **<0.01** |
|  | STX1B | Syntaxin 1B | –0.17 ± 0.15 | –0.25 | 35.01 | 0.02 | >0.01 |
| RNA metabolic process | CPSF4 | Cleavage and polyadenylation specific factor 4 | –0.41 ± 0.16 | –0.25 | 34.27 | 0.23 | **<0.01** |
|  | CRNKL1 | Crooked neck pre-mRNA splicing factor 1 | –0.04 ± 0.17 | –0.25 | 34.91 | 0.04 | >0.01 |
|  | DDX23 | DEAD (Asp-Glu-Ala-Asp) box polypeptide 23 | –0.36 ± 0.35 | –0.25 | 32.22 | 0.01 | >0.01 |
|  | PRPF8 | Pre-mRNA processing factor 8 | –0.33 ± 0.08 | –0.25 | 33.57 | 0.26 | **<0.01** |
|  | WDR33 | WD repeat domain 33 | –0.29 ± 0.12 | –0.25 | 33.77 | 0.14 | **<0.01** |
